# Evaluating Flow-Focused Microfluidic Device Fabrication Techniques for Silk Fibroin Microgel Production

**DOI:** 10.1101/2025.02.02.636143

**Authors:** Mahsa Haghighattalab, Fatemeh Karimi, Nona Farbehi, Jelena Rnjak-Kovacina

## Abstract

Microgels, or micro-scale hydrogels, are versatile emerging biomaterials. They absorb large amounts of water and facilitate molecule transfer. Their tunable size and shape and the capacity to form 3D scaffolds with microscale porosity offer significant advantages over traditional bulk hydrogels. Various strategies exist for fabricating microgels with microfluidic techniques offering the most control of microgel properties. This study compares three microfluidic device fabrication techniques—maskless photolithography, laser engraving, and 3D printing—for making photo-crosslinked silk fibroin microgels. Silk microgels were fabricated via water-in-oil microfluidics and crosslinked through di-tyrosine bonds between native tyrosine residues in silk. Microfluidic devices with target channel depths of 50, 100, or 400 µm were successfully fabricated, with each technique offering unique advantages and limitations. Maskless photolithography provided the highest channel patterning accuracy and smoothest profiles but was costly and required specialized facilities. Laser engraving was affordable but labor-intensive, with lower accuracy in the X-Y plane for deeper channels. 3D printing was user-friendly and affordable but resulted in low accuracy for 50 µm channels and rougher channel profiles, leading to larger microgels and less efficient microgel formation than the other techniques. Spherical silk microgels with diameters between 50 µm and > 400 µm were produced by modulating channel depth and flow rates, while rod-shaped microgels were made by crosslinking in the outlet tubing. Silk molecular weight was adjusted to control surface porosity and compressive modulus, with lower molecular weight silk resulting in higher porosity and softer microgels (modulated between ∼ 40 kPa and ∼ 590 kPa) while maintaining consistent size.

This study demonstrates the versatility and effectiveness of different microfluidic device fabrication techniques for producing photo-crosslinked silk microgels with tunable properties for biomedical applications.

## 1. Introduction

Hydrogels are polymeric networks that absorb large amounts of water, exhibiting swelling behavior that facilitates the transfer of small and large molecules [1]. Hydrogels are among the most popular biomaterials, with diverse biomedical applications such as tissue engineering and drug delivery [2–4]. In recent decades, significant effort has been made to advance hydrogel structures to introduce spatiotemporal gradients of physiochemical properties toward building more functional and biomimetic scaffolds [5,6]. Despite their beneficial properties and potential, the application of traditional bulk hydrogels is limited by their lack of micro-scale porosity, which restricts cell and blood vessels invasion, leading to fibrous encapsulation when implanted in the body [7]. Moreover, traditional bulk hydrogels lack modularity, essential for mimicking complex and dynamic biological systems.

Microgels are micro-scale hydrogels that have gained popularity in recent years [8]. They offer similar advantages and applications as hydrogels, including preserving the bioactivity of molecules [9], drug delivery [10], cells expansion [11], and tissue regeneration [12]., but their micro-scale offers unique advantages. These microgels can be annealed together to form microgel scaffolds which are networks of interconnected microgel building blocks that collectively form a porous and modular hydrogel matrix, essential for effective tissue remodeling. [13]. Various strategies exist for fabricating microgels, including mechanical fragmentation, bulk water-in-oil emulsion, and microfluidic techniques [14–17]. Microfluidics, in particular, is valuable for the precise control of the size [9], shape [18], and the structure of microgels [19]. Various fabrication techniques, such as photolithography, laser engraving, and 3D printing, can be used to create microfluidic devices, depending on the required feature size and the specific application needs.

Photolithography is a high-end technology that offers the highest feature resolution, reaching submicrometer scales [20,21]. This technique involves creating microchannel patterns using a photoresist on a silicon wafer, followed by casting a material (typically polydimethylsiloxane (PDMS)) to create the final pattern of the microfluidic device. Although only a single-step (excluding post-hydrophobicity treatment) molding is needed to make PDMS-based microfluidic devices, the master fabrication process is relatively costly and complex. Two other affordable and readily available fabrication options are CO_2_ laser-assisted acrylic engraving [22] and 3D printing [23]. Laser engraving can be used to create microchannel patterns on thermoplastic acrylic sheets, such as Poly(Methyl MethAcrylate) (PMMA) by engraving with CO_2_ lasers [24]. If the engraved microchannels are negative, two replication steps are needed to fabricate a PDMS device with negative channels [25]. Although this process is affordable, quick, and straightforward, it is time-consuming and labor-intensive due to the manual optimization of the fabrication parameters and the limited reusability of the sacrificial mold due to shrinkage and deformation. Another conventional microfluidics fabrication technique is 3D printing, where a master or final device is created to form microchannels [26,27]. While 3D printing offers significant advantages in fabricating microfluidic devices, certain challenges persist. Notably, some 3D printing materials have been found to exhibit cytotoxicity, limiting their biocompatibility in cell-based assays. Additionally, the relatively rough surface finish of 3D-printed microfluidic master molds can lead to higher surface roughness of the channels which results in inconsistencies in droplet formation, negatively affecting the reproducibility and precision of experiments [28]. Additional post-processing steps, such as solvent treatment, are often necessary to smooth surfaces and improve the reliability of device functionality. These steps are crucial for optimizing 3D-printed materials used in microfluidic applications.

This work compares the three common microfluidic device fabrication techniques-maskless photolithography, laser engraving, and 3D printing for making silk fibroin microgels. Silk fibroin (henceforth referred to as ‘silk’) is an abundantly available natural polymer widely explored for a range of biomedical applications, such as tissue engineering, drug delivery, and biosensing due to its cytocompatibility and highly tunable mechanical, morphological, and degradation profiles [29]. Silk microgels have been fabricated using a range of techniques, including dripping solvent coagulation [30], dripping water-in-oil emulsion [31], and microfluidics [32–34]. In most of these studies, silk microgels are stabilized via physical crosslinking involving the native β-sheet secondary protein structures in silk. However, β-sheets can result in brittle materials, and recently, covalent di-tyrosine crosslinking has become a popular means of producing elastomeric silk hydrogels. The first di-tyrosine crosslinked silk microgels were made using visible light-based photocrosslinking in the presence of the photo-initiator ruthenium(II)hexahydrate (Ru(II)(boy)32+) (Ru) and sodium persulphate (SPS) as the electron acceptor [25]. This method is advantageous as it stabilizes silk without chemical modification; it is also rapid, cytocompatible, and supports the functionalization of silk with bioactive molecules.

The microfluidic devices used for fabricating di-tyrosine crosslinked silk microgels were fabricated using laser engraving to generate microgel populations with average diameters of ∼ 100 µm or ∼ 400 µm. However, the channel features were not optimized in this study, and the effect of channel features on photocrosslinked silk microgel properties remains unknown. In this work, we investigated the feasibility of three different microfluidic device fabrication methods to generate channel depths of 50, 100, and 400 µm, as different channel depths result in different droplet sizes at the same fluid flow rate. We compared the final PDMS devices in terms of patterning accuracy, reproducibility, and channel surface roughness, which are crucial for droplet generation and microgel formation. The efficacy of these techniques was assessed in the context of silk droplet formation and microgel generation. Moreover, we presented and critically evaluated the in-process parameters for generating silk microgels with target micro-scale features such as size and shape. In addition to regulating the morphological properties at the micro-scale, we also showed how inherent properties such as the nano-scale porosity and compressive elastic modulus can be tuned for different applications. Tuning these micro- and nano-scale properties is important to tailor biomaterials to better mimic the complexity of biological systems.

## 2. Materials and methods

### 2.1 Materials

*Bombyx mori* silk cocoons were supplied by Sato Yama, Japan. Lithium bromide (LiBr), Sodium carbonate (Na2CO3), Tris(2,2′-bipyridyl)dichlororuthenium(II) hexahydrate, Sodium persulfate (SPS), heavy mineral oil, Span® 80, and Trichloro(1*H*,1*H*,2*H*,2*H*-perfluorooctyl) silane were purchased from Sigma-Aldrich (Merck, UK). Dialysis tubing was supplied from Thermo Fisher (3,500 MWCO, SnakeSkin^𝑇𝑀^, USA). Masterflex microbore transfer tubing (0.020” ID x 0.060” OD) was used as microfluidic tubing. Polydimethylsiloxane (PDMS) was SYLGARD™ 184 silicone elastomer kit. FUSION 4000-X dual independent channels syringe pump was used to control the fluids flow, CHEMYX 30W portable LED worklight was used to photocrosslink silk microgels, and Femto Diener plasma surface treatment machine was used to oxygen plasma treat the substrates.

### 2.2 Microfluidic device design

The microfluidic device channel geometry was designed based on Karimi et al.’s previous publication [25] and is illustrated in detail in Figure S1 (Supporting Information).

### 2.3 CO_2_ laser-assisted engraving of acrylic masters

Transparent acrylic sheets (4.5 mm thickness) were used as master substrates and engraved using a CO_2_ laser cutter (Speedy 360) machine. Different powers ranging from 10% to 100% were used to engrave acrylic substrates (n = 3-5) to obtain various channel depths. The power needed to achieve three targeted depths of 50, 100, and 400 µm was estimated using fitting a linear regression model and interpolation (Figure S2). The in-process parameters are summarized in Table 1.

**Table 1.** In-process parameters used for laser engraving of acrylic substrates. PPI = pulses per inch.

| Process | Power (%) | Speed (%) | PPI/Hz | Passes | Air Assist | Direction | Z-Offset |
| --- | --- | --- | --- | --- | --- | --- | --- |
| Engrave | 10-100 | 30 | 500 PPI | 1 | On | Top down | 0 |
| Cut | 95 | 0.5 | 1000 Hz | 1 | On | - | 0 |

### 2.4 Fabrication of the microfluidic device from laser-engraved substrates

To fabricate the final PDMS microfluidic device, two casting steps were required to replicate the negative channel patterns from the engraved substrate to the final PDMS substrate with negative channels (Figure 1A (i) (dxf file attached)). The first casting step involved transferring the negative acrylic channel patterns to positive PDMS channels, creating a sacrificial mold. The laser-engraved acrylic master was gently cleaned with an isopropyl wipe and air-dried to remove dust and debris. PDMS was used at a 10:1 ratio of base to crosslinker, followed by degassing in a vacuum chamber before overnight curing at 37°C. The back of the acrylic substrate was adhered to a silicon rubber mold using double-sided tape, followed by casting 10 g of PDMS. After demolding, the sacrificial mold was treated with oxygen plasma (1 min, at the power of 50 W and 𝑂_2_ flow rate of 5 sccm) and silanized. Silanization was done by 3h of silane vapor deposition by placing 20 µl of (Trichloro(1H,1H,2H,2H-perfluorooctyl)) (silane) in a desiccator and vacuuming under fume hood. In the second casting step, 10 g of PDMS was cast into the sacrificial mold to transfer the positive to negative channel patterns, forming the final substrate. After demolding the PDMS substrate with the negative channel pattern, inlets, and outlet holes were created by vertically punching the substrate using a 1.2 mm biopsy puncher (ProSciTech). The PDMS substrate and a glass slide were cleaned with an isopropyl wipe, and 70% ethanol, respectively, followed by air drying. To seal the device channels, both the PDMS substrate and glass slide were treated with oxygen plasma and bonded together. To improve the hydrophobicity of the channels, channels were filled with RainX solution for 20 minutes, aspirated, and dried by air. The inlets and outlets were kept covered using tape until the device was used.

**Figure 1.**
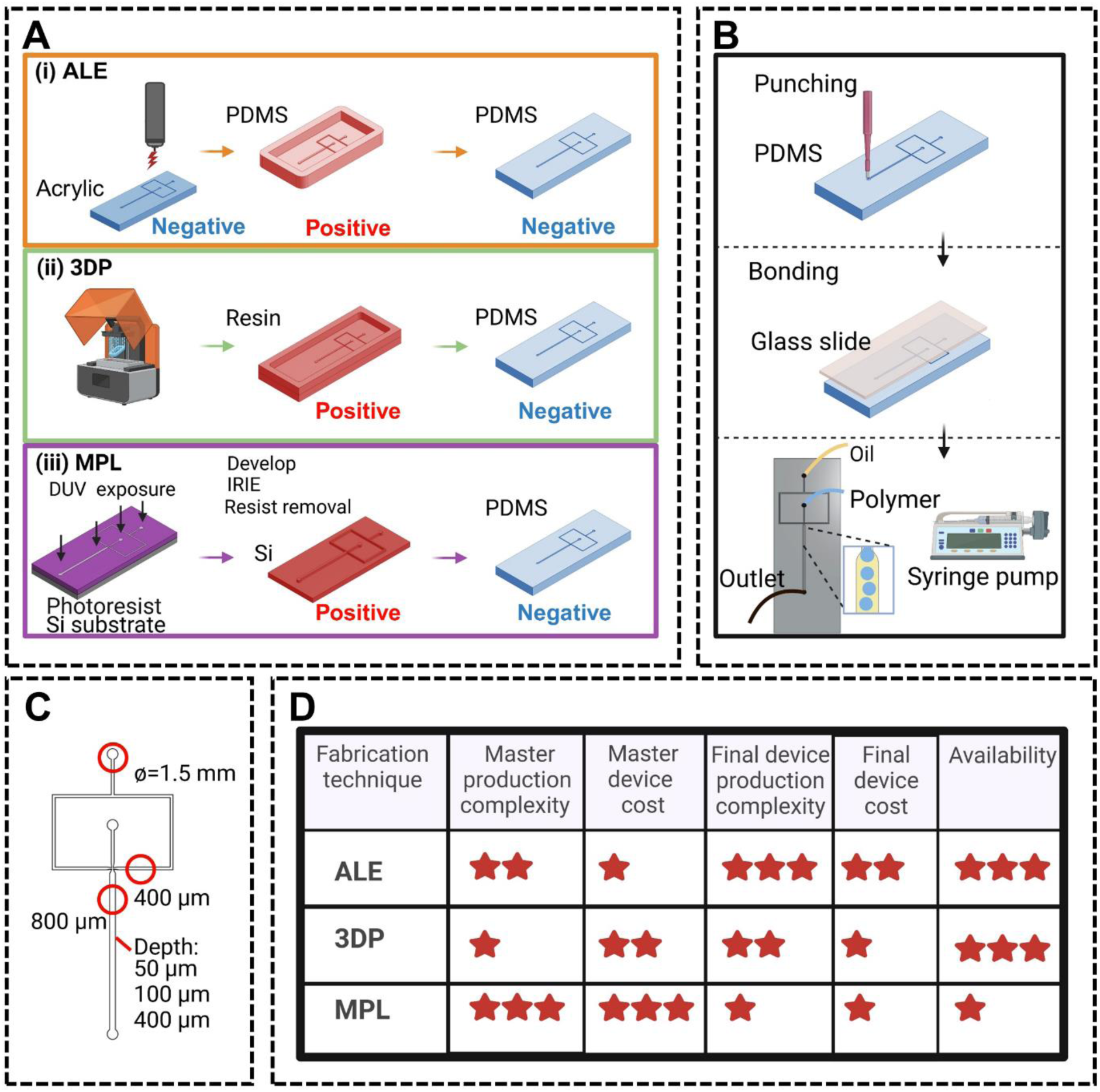
Microfluidic device fabrication by acrylic laser engraving (ALE), 3D printing (3DP), and maskless photolithography (MPL). A) An overview of the master device fabrication steps using i) ALE, ii) 3DP, and iii) MPL. B) Fabrication steps from a master device to the final PDMS microfluidic device. C) Microfluidic device design used in this study. A detailed design drawing is available in Figure S1. D) Comparison of the different microfluidic fabrication techniques used in this study.

### 2.5 Fabrication of microfluidic devices using 3D printing

3D printed master molds with positive channel patterns were made of biomedical-grade clear resin (Formlabs Clear Resin) using a SLA printer (Formlab, Form 3B+) (Figure 1A (ii), STL files attached). The supports were printed at the back of the molds at a 45° angle. After printing, the 3D printed molds were post-processed by soaking in isopropyl alcohol for 20-30 minutes and sonicated for 4-5 minutes to remove any uncured material. The supports were then removed from the back of the resin molds and sanded to flatten the surface. To make the PDMS microfluidic device from the 3D printed masters, the resin molds were cleaned with isopropyl wipe, air-dried, oxygen plasma treated, and silanized (as described above). 10g of PDMS (10:1 ratio of base and curing agent) was poured into the resin mold followed by degassing in a vacuum chamber and cured at 60 °C for 3-4 hours. The PDMS substrate was then demolded using a scalpel and spatula and punched vertically to create the inlets and outlet. Punched PDMS substrate and a glass slide were cleaned with an isopropyl wipe, and ethanol (70%), respectively, then air-dried, oxygen plasma treated and bonded together. RainX solution was used for 20 minutes to decrease the channels’ hydrophilicity caused by plasma treatment. RainX aspirated and microfluidic devices allowed to dry by air overnight. Microfluidic device inlets and outlet were kept covered using tape until the usage.

### 2.6 Fabrication of microfluidic devices using maskless photolithography

Silicon wafers with positive channels were made using maskless photolithography at the UNSW Node of the Australian National Fabrication Facility (Figure 1A (iii), dxf and dwg files attached). Briefly, AZ ECI 3012 positive photoresist was spun on the Si wafer and exposed with the direct write tool (maskless photolithography). The areas were then etched down with ICP-RIE (Bosch Process), and the resist removed. The silicon wafer with patterns was placed in a petri dish, oxygen plasma treated, and silanized (as described above). PDMS poured onto silicon wafer with patterns, followed by degassing in a vacuum chamber and curing at 60°C for 3-4 hours. The PDMS substrate was then demolded using a scalpel and spatula and punched vertically to create the inlets and outlet. Finally, PDMS substrates and glass slide were cleaned with isopropyl wipe, and 70% ethanol, respectively, air-dried, and bonded together using plasma treatment (as described above). Then, channels filled with RainX and aspirated after 20 min and microfluidic devices allowed to dry by air overnight. Microfluidic device inlets and outlet were kept covered using tape until the usage.

### 2.7 Characterization of the channel profiles

The channel profiles were assessed using an optical profilometer (ContourGT, Bruker). The depth of the channels was measured as the peak-to-valley height value (Figure S3A). The width of the channels was measured from images acquired by an optical microscope (Leica) and ImageJ software, or an optical profilometer (Figure S3B). The size variation (%) and accuracy (%) of geometry patterning in X and Z-axes were calculated using equations 1 and 2, respectively. The accuracy of channel geometry patterning measures how close the pattern geometry is to the nominal value (the CAD file dimensions). To consider the effect of feature size on accuracy, the accuracy of geometry patterning was calculated for both the smallest and biggest widths in the design, 400 µm and 800 µm.

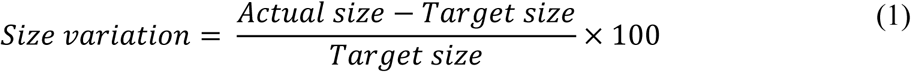

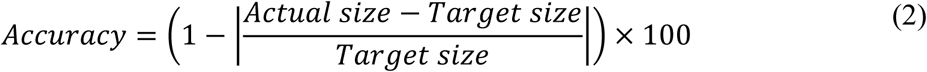

Surface index (SI) was measured using a Bruker optical profilometer to indicate surface morphology or roughness. SI is the measure of surface area (effective area) in the lateral resolution over a perfectly flat area (geometric area). Thus, the higher the SI, the rougher the surface (Figure S3C) [35].

Measurements were done on the final PDMS replica substrates (unless stated otherwise) and expressed as the average of measures over three regions in each substrate, with n = 3-5 prototyping replications.

### 2.8 Isolation of silk fibroin from silk cocoons

Silk fibroin solution was made based on Rockwood and colleagues’ protocol [29]. Briefly, *Bombyx mori* silk cocoons were cut into small pieces, followed by boiling in 0.02M Na2Co3 (degumming, 10 min, 30 min, or 60 min) to extract and remove the glue-like sericin protein. Degummed silk fibers were rinsed with deionized water and dried overnight. Dry fibers were pulled apart to increase the surface area and incubated in 9.3M LiBr (0.25g fibers/mL for 30 min, and 60 min degummed fibers, and 0.2g fibers/ml for 10 min degummed fibers) at 60°C for 4 h. LiBr was removed by dialysis against MilliQ water in Snakeskin tubing (3,500 MWCO) for 3 days. Regenerated silk fibroin solution was collected and centrifuged two times at 8700 rpm for 15 minutes at 4°C. The solution was stored at 4°C until required. To determine the weight-to-volume percentage (%w/v), 500 µl of silk solution (n = 3) was incubated at 60°C in a weigh boat overnight to dry, and then %w/v was calculated using equation 6.

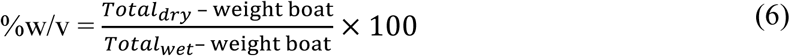

### 2.9 Fabrication of photocrosslinked silk fibroin microgels

Microgels were made in the microfluidic devices fabricated using different fabrication techniques described above. Each device had two inlets, one for heavy mineral oil supplemented with 6% Span-80 and the other for the silk microgel precursor solution. The microgel pre-cursor solution consisted of 3% w/v silk solution (10 min, 30 min, or 60 min degummed), 0.25 mM Tris2,2′bipyridyl)dichlororuthenium(II) hexahydrate (Ru), and 2.5 mM sodium persulfate (SPS).

After collecting silk microgels/droplets in a tube containing 2-3 ml oil phase, microgels/droplets were photocrosslinked in the presence of visible light for 20 min to stabilize. Then, microgels immersed in the oil phase were centrifuged at 1000 rpm for 3 min, and the oil was discarded as much as possible. In the following, 4-5 ml 1X phosphate buffered saline (PBS) were added to microgels, shaken to dispatch agglomerated microgels, filled with PBS, and centrifuged at 3000 rpm for 5 min. The supernatant was discarded, and the tube wall was cleaned using a rolled Kim wipe (2 times). Again, 3-4 ml 1X PBS was added, and the tube was changed. Depending on the size of the microgels, this process was repeated 2-3 times to wash the oil out completely.

For washing silk microgels smaller than 100 µm (diameter), after taking the oil out, microgels were washed with a 1% Tween 20 washing agent (nonionic detergent). This was followed by washing in 1X PBS for 2-3 times.

### 2.10 Quantification of silk droplet generation rate

The droplet formation of 3% w/v silk solution was captured using a high-speed camera (VEO 410L, Phantom) mounted on an optical microscope (CKX53, Olympus). The droplet generation images were captured at either 60 fps or 100 fps, depending on the speed of generation. The number of droplets formed per second was counted over 3 to 5 seconds for each condition.

### 2.11 Silk microgels size and quantification

To quantify the microgel size, images were captured using an optical microscope (CKX53, Olympus) (Figure S4), and individual microgel area was measured using ImageJ software (NIH). The diameter was then calculated from the surface area value (n = 100).

### 2.12 Quantification of the microgel surface porosity

Silk fibroin microgels were packed via centrifugation at 3000 rpm (5810 R, Eppendorf) for 5 min in Eppendorf tube 1.5 ml (Figure 6C). Packed microgels were sequentially dehydrated via two washes in each ethanol (30, 50, 70, 80, 90, 95, and 100 (3 X) v/v%) using a Biowave Pro+ microwave processor (Pelco). The dehydrated samples were dried using an EM CPD300 critical point dryer (Leica), mounted on a stub using double-sided carbon tape, and coated with 15 nm platinum using a Q300T D Plus sputter (Quorum). FESEM (field-emission scanning electron microscope) images were taken using Nova NanoSEM 230 machine under a high vacuum using a beam power of 5 kV, and probe current of 30 μA. To assess the surface porosity (%) of silk microgels, 5 images of each condition at 20000x magnification were analyzed. 2D average void space fraction was measured using ImageJ software (NIH) and expressed as the surface porosity (%).

### 2.13 Confocal microscope imaging of silk microgels

Silk fibroin microgels were stained with Tetramethylrhodamine (TRITC) to capture the confocal images. 50 μg/ml TRITC was added to silk microgels at a 2:1 volume ratio and incubated overnight at 4 °C. Microgels were then washed in 1X PBS 2-3 times before imaging [25]. Confocal images were taken using a confocal laser scanning microscope (Olympus FluoView FV3000) in variable beam focus (VBF) mode (detection wavelength of 570 - 620 nm) and laser wavelength of 561 nm.

### 2.14 Analysis of the silk solution viscosity

The viscosity of solutions was measured using a Brookfield DV2T viscometer. The viscosity of the silk solutions with different degumming times of 10, 30, and 60 minutes was measured at three different temperatures of 4 °C, 25 °C, and 37 °C (n > 5).

### 2.15 Mechanical testing of silk microgels

The mechanical properties of single microgels were studied using the MicroTester LT instrument (CellScale). Single microgel deformation was assessed using parallel plate compression testing, in which a platen (3×3 mm) compressed the microgel (placed on an anvil) up to 100 µm displacement magnitude in 20 seconds. Depending on the maximum force, two different beam diameters were used (0.4064, and 0.5588). The Hertzian half-space contact mechanics model was used to calculate the compressive elastic modulus (E) (Figure S5A) [36]. This model can be used for small deformations (𝜀 ≤ 10%, **equation 7**) of soft biomaterials with the assumption of Poisson’s ratio (ѵ) to be 0.5, and neglection of the reaction force (-F). Then, elasticity was calculated from displacement (δ), the radius of the sphere (R), and force (F) at each time point using **equation 8**.

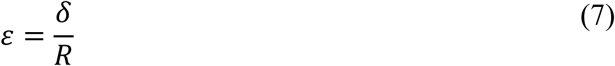

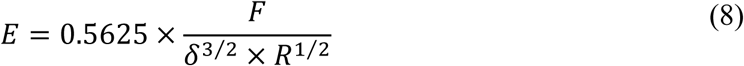

### 2.16 Statistical analyses

Data are presented as mean ± standard deviation (SD). Statistically significant differences were determined by student t-test or one-way analysis of variance (ANOVA) and the Tukey post-test using GraphPad Prism 9 (GraphPad San Diego, CA, USA). Statistical significance was reported at p < 0.05 and indicated in the figures as * p < 0.05, ** p < 0.01, *** p < 0.001, and **** p < 0.0001, or ns (not significant).

## 3. Results and Discussion

### 3.1. Fabrication of flow-focusing microfluidic devices

Microfluidic devices were fabricated using three techniques-laser engraving, 3D printing, and maskless photolithography (**Figure 1A**). The devices consisted of two fluid inlet ports, flow-focusing channels (target width of 400 µm in inlets, 800 µm in the outlet, and channel depths of 50 µm, 100 µm, or 400 µm), and one outlet port (**Figure 1B**, detailed dimensions **Figure S1**). PDMS was cast in the mold to create the negative channels, and inlets and outlets were generated using a biopsy punch (ID: 1.2 mm, and OD:1.5mm). The PDMS was then bonded to a glass slide using oxygen plasma treatment, and tubing was connected to inlets and the outlet to complete the microfluidic devices for microgel fabrication (**Figure 1C**). The advantages and disadvantages of each fabrication method, based on equipment availability and user workflows, are outlined in **Figure 1D**.

Laser engraving requires manual optimization of the laser power to reach the desired channel depth. The laser power was adjusted between 10% and 100%, which resulted in channels with depths ranging from 1.41 ± 0.14 µm to 663.67 ± 5.79 µm (**Figure S2A**). Target channel depths of 50, 100, and 400 µm were made in raster mode with a laser power of 22.5%, 32.5%, and 80%, respectively. The direction of the laser scanning was also a crucial consideration, as horizontal scanning resulted in an undesirable scalloped pattern (**Figure S2B**). Laser engraving was the most accessible and cheapest fabrication method for making the master substrate. However, laser engraving was based on creating negative channel patterns, which was labor intensive considering the two steps of PDMS casting needed to transform the negative patterns from the acrylic substrate to the negative PDMS substrate.

3D printing was the easiest method for generating the master mold, while maskless photolithography involved a complex multistep process requiring specialized fabrication facilities. Both methods resulted in positive channel patterns, so they were easier and cheaper for the fabrication of the final PDMS device.

### 3.2. Microfluidic device channel profile characteristics

The accuracy of each fabrication technique was assessed by analyzing the channel profiles of the final PDMS replica devices. To verify that the PDMS devices are accurate replicas of the master molds, the channel depth, width, and surface topography (expressed as surface index) of the master substrates and PDMS device fabricated via laser engraving were compared. Laser engraving was chosen because it involves multiple molding steps, making it more prone to inaccuracies than other techniques. The PDMS devices were found to be accurate replicas of the master molds, with no significant differences in any of the assessed parameters, regardless of the target channel depth (**Figure 2**). This ensures that the final PDMS replica retains the same microchannel feature size and properties as the original master substrate.

**Figure 2.**
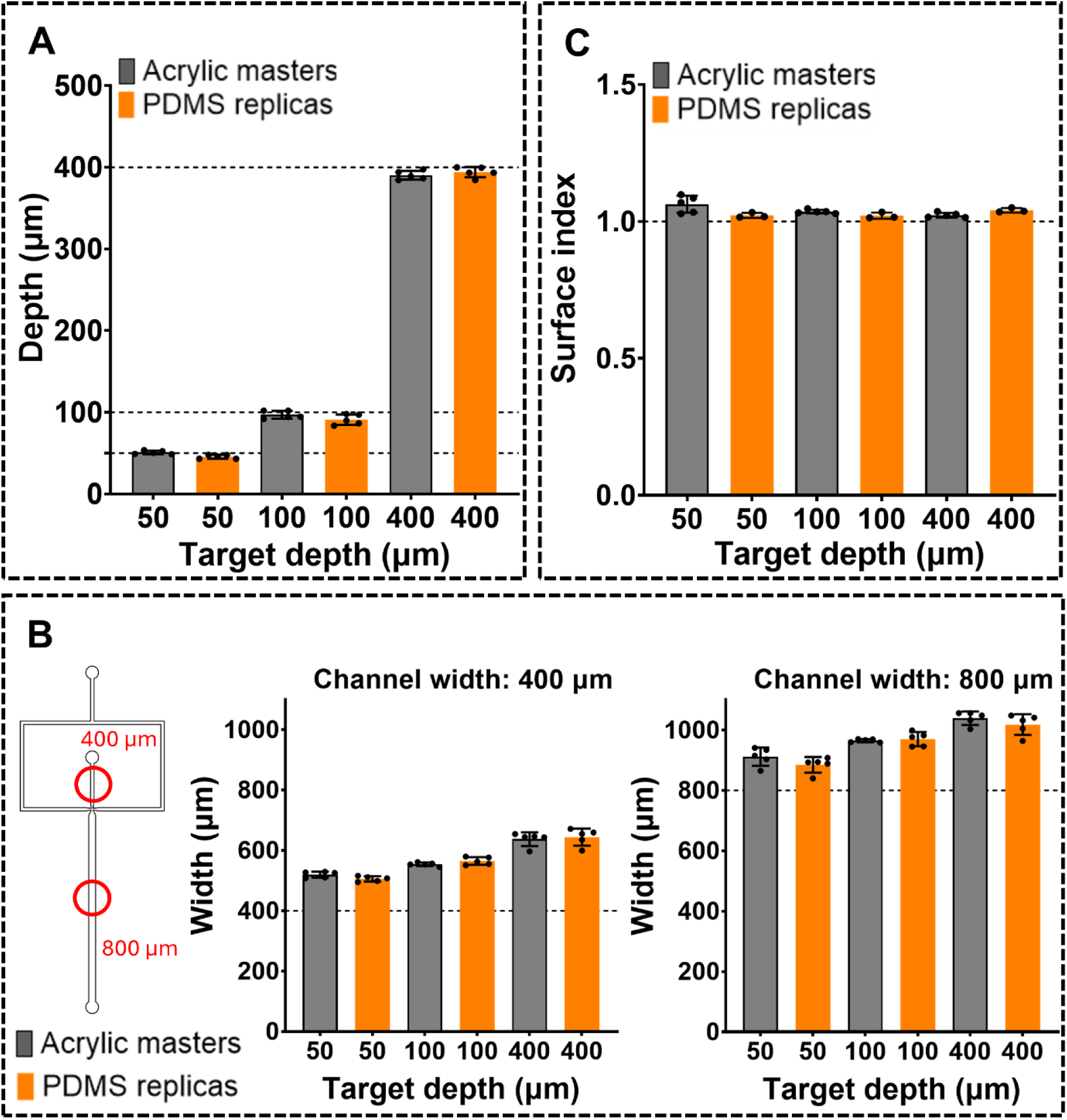
Comparison of the channel profiles of the acrylic master with the final PDMS devices. **A)** Microfluidic channel depths at 50 µm, 100 µm, and 400 µm target depth. **B)** Microfluidic channel widths at 400 µm (inlet) and 800 µm (outlet) target widths **C)** Surface index of the channels as an indicator of surface roughness. Dashed lines in each panel indicate the target value. Data are mean ± SD, N = 3-5.

Having established that the PDMS devices are accurate replicas of the master molds, the channel profile of the final PDMS devices generated via each fabrication technique was assessed using an optical profilometer, as shown in **Figure 3**. The accuracy of the channel depth was dependent both on the fabrication technique and the target depth of the channel (**Figure 3B, C**). For the 50 µm target channel depth, 3D printing resulted in significantly deeper channels (p < 0.0001) than the target, with only 63% accuracy, while the other two techniques were > 90% accurate (**Figure 3B, C**). For the 100 µm and 400 µm target channel depths, all techniques achieved good accuracy of > 90%, but maskless photolithography proved to be the most accurate (**Figure 3B, C**).

**Figure 3.**
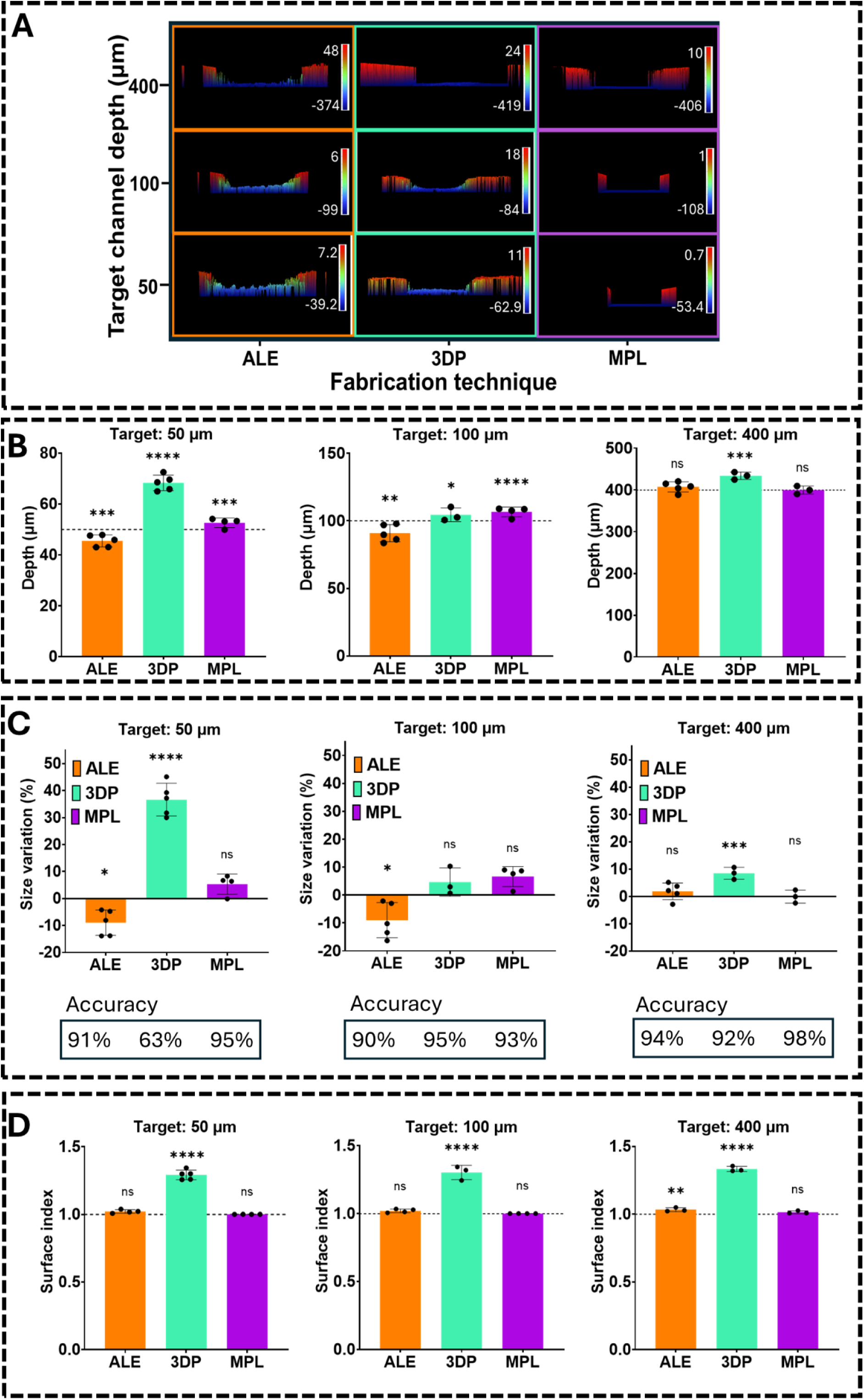
Channel profiles of the final PDMS microfluidic devices made using acrylic laser engraving (ALE), 3D printing (3DP), or maskless photolithography (MPL). **A)** Representative microfluidic channel profiles assessed by optical profilometry. **B)** Channel depths for different target depth values (50, 100, or 400 µm). **C)** Size variation (%) and accuracy of channel depth patterning. **D)** Surface index of microfluidic channels. Dashed lines indicate target values. Data are mean ± SD, N = 3-5. * p < 0.05, ** p < 0.01, *** p < 0.001, **** p < 0.0001, relative to target value.

Surface index was measured to indicate surface roughness, which can potentially affect the fluid flow dynamics in microchannels (**Figure 3D**). The surface index of PDMS replicas from the maskless photolithography method was the lowest among all the techniques. Surface index was measured at 1.002 ± 0.001, 1.001 ± 0.000, and 1.015 ± 0.009 for 50 µm, 100 µm, and 400 µm target channel depths, respectively. These measures show that the channel profiles were close to perfectly smooth (true value of 1). In contrast, the roughest surface was in PDMS replicas from the 3D printing method with the highest surface index value of 1.293 ± 0.032, 1.303 ± 0.042, and 1.385 ± 0.007 for 50 µm, 100 µm, and 400 µm target channel depths. Across all techniques, no significant difference was observed in the surface index between different channel depths, indicating that the surface roughness is independent of the channel depth.

### 3.3. Characterization of the microfluidic device channel pattern in the X-Y plane

The microfluidic channel widths were assessed to compare the dimensional variation and accuracy across different techniques when patterning 400 µm and 800 µm channel widths (**Figure 4**). Photographs of the channels generated via the different techniques showed clear channel contours and edges in maskless photolithography and more diffuse edges and less well-replicated channel contours in the other two techniques (**Figure 4A**).

**Figure 4.**
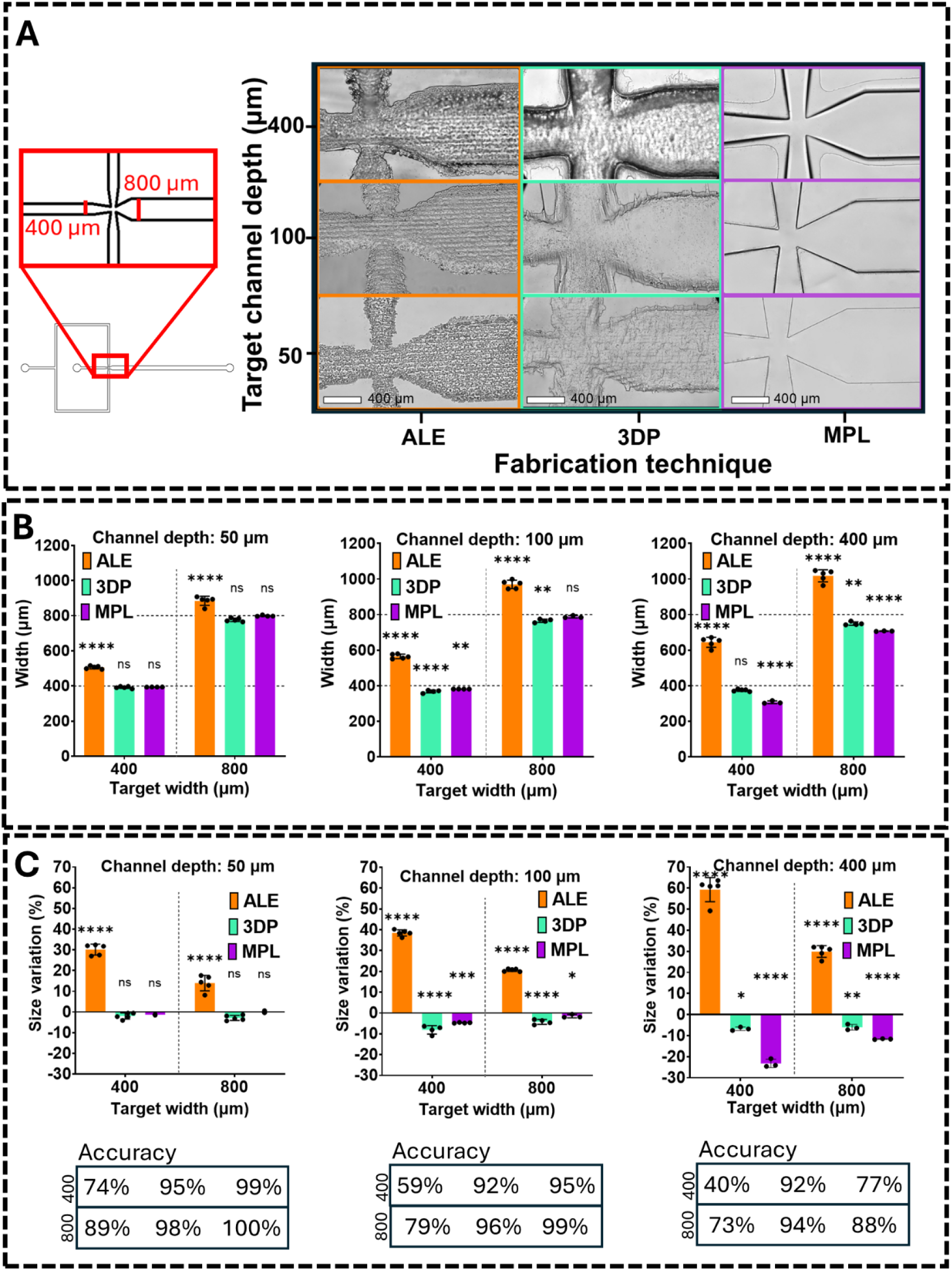
Analysis of the width of channels made using acrylic laser engraving (ALE), 3D printing (3DP), or maskless photolithography (MPL). **A)** Microscopic images of the final microfluidic devices’ T junction made using different fabrication techniques. **B)** Microfluidic channel widths. **C)** Size variation (%) and accuracy of channel width patterning. Data are mean ± SD, N = 3-5. * p < 0.05, ** p < 0.01, *** p < 0.001, **** p < 0.0001, relative to target value.

Acrylic laser engraving exhibited the greatest width variation under all conditions, with channel width accuracy as low as 40% when fabricating 400 µm wide channels (**Figure 4B, C**). The channels fabricated using this method were consistently wider than the target width. Interestingly, the accuracy of channel geometry patterning using this technique decreased with the increase in laser power (engraved depth) (**Figure 4B**). This variability is likely due to the nature of the laser engraving process, which involves burning the material using laser power. It is possible that with further optimization of the laser speed and power, this accuracy may be improved, as it was initially chosen based on the target channel depth.

3D printing and maskless photolithography resulted in the more accurate fabrication of the target channel widths, with > 90% accuracy across all conditions, except notably for maskless photolithography in the 400 µm channel depth, where the accuracy was 77% for 400 µm and 88% for 800 µm channel width (**Figure 4C**). Interestingly, unlike for channel depth, 3D printing achieved excellent accuracy in the X-Y plane, with accuracy > 92% in all conditions (**Figure 4B, C**). This is likely due to the nature of the SLA process, where the X-Y plane accuracy is primarily determined by the precision of the laser or projector used to cure the resin and can achieve fine resolution, while the Z-plane accuracy is determined by the movement of the build platform to achieve a specific layer height, potentially introducing inaccuracies [37]. 3D printing and maskless photolithography consistently resulted in channels of target width or slightly narrower, unlike laser engraving, which consistently resulted in wider than target channels. Overall, the MLP technique had the highest accuracy in patterning the channel geometry in the X-Y plane, followed by 3D printing and laser etching.

### 3.4. The effect of microfluidic device fabrication techniques on silk microgels

Microfluidics droplet formation in passive mode depends on multiple parameters, including channel geometry, fluid flow rates, and fluid properties [38–40]. Since different microfluidic device fabrication techniques resulted in channels with different properties, silk microgels were produced in devices fabricated via each technique using set flow conditions (0.2 flow rate ratio 𝑄_𝑆𝑖𝑙𝑘_: 𝑄_𝑂𝑖𝑙_, 𝑄_𝑂𝑖𝑙_ = 500 µl/h) to assess the effect of the channel properties on silk microgel size, shape, and surface porosity. **Figure 5A** illustrates silk droplet formation in microfluidic devices, with a clear effect of device fabrication technique and channel depth on droplet formation profiles. Deeper channels resulted in larger droplets and a slower droplet generation rate, as evidenced by single droplet formation and clear separation between individual droplets in the 400 µm channels, regardless of the fabrication method. In contrast, shallower channels produced smaller droplets at a faster rate, with droplets flowing in multiple streams and close to each other. Interestingly, 3D printed microfluidic devices exhibited notably different droplet generation profiles compared to other techniques. This was confirmed by quantifying the droplet generation rate (**Figure 5B**) in a 100 µm channel device at different flow rates. Regardless of the flow rate, droplets were generated significantly slower in the 3D printed devices. Given that the accuracy of the 3D printing technique for 100 µm channels was comparable to the other techniques in terms of channel depth and width (**Figure 3, 4**), it is likely that the significantly rougher surface of the 3D printed channels affected droplet formation by causing energy loss and velocity reduction.

**Figure 5.**
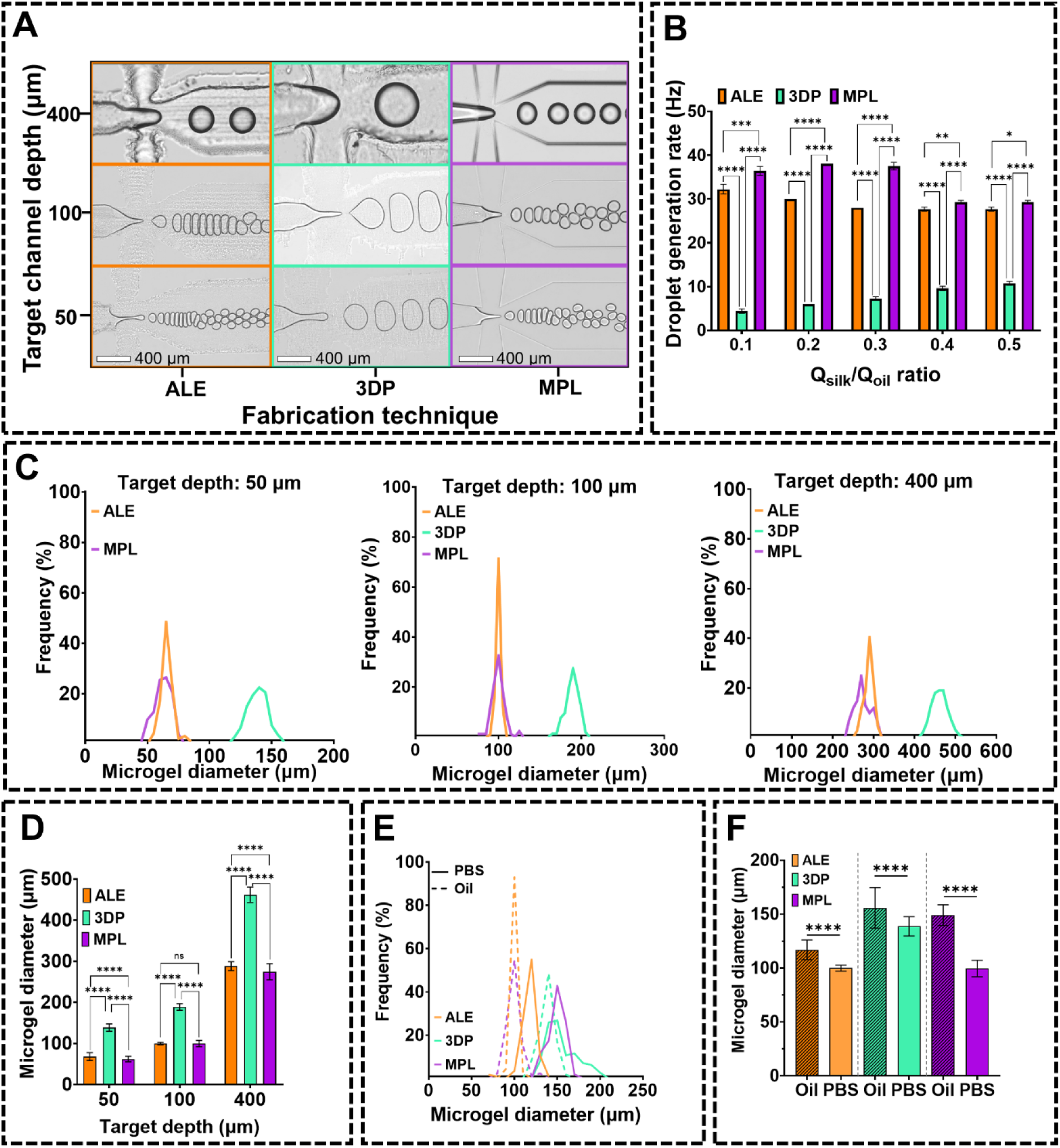
The effect of different microfluidic device fabrication techniques on silk microgels formation. **A)** Droplet formation of silk fibroin solution at a constant flow rate ratio of 0.2 (𝑄_𝑂𝑖𝑙_ = 500 𝜇𝑙/ℎ𝑟). **B)** Droplet generation rate in different fabrication techniques (target channel depth of 100 μm). **C)** Frequency distribution of microgel diameter made by a constant flow rate ratio of 0.2 (𝑄_𝑂𝑖𝑙_ = 500 𝜇𝑙⁄ℎ𝑟) using different device fabrication techniques. Microgel size was quantified after washing in PBS. **D)** Diameter of silk microgels made with a constant flow rate ratio of 0.2 (𝑄_𝑂𝑖𝑙_ = 500 𝜇𝑙⁄ℎ𝑟). Microgel size was quantified after washing in PBS. **E)** Frequency distribution of silk microgels diameter made with a target channel size of 100 µm at a constant flow rate ratio of 0.2 (𝑄_𝑂𝑖𝑙_ = 500 𝜇𝑙⁄ℎ𝑟) showing microgels in oil and following washing in PBS. **F)** Silk microgels diameter made with a target channel size of 100 µm at a constant flow rate ratio of 0.2 (𝑄_𝑂𝑖𝑙_ = 500 𝜇𝑙⁄ℎ𝑟) showing microgels in oil and following washing in PBS. ALE; acrylic laser etching, 3DP; 3D printing, MPL; maskless photolithography. Data are mean ± SD, N = 100, * p < 0.05, ** p < 0.01, *** p < 0.001, **** p < 0.0001, relative to target value.

The microgel diameter was polydisperse regardless of the device fabrication technique. However, microgels generated in the acrylic laser-etched devices had the narrowest diameter distribution profiles, especially in 100 µm channels (**Figure 5C**). The microgel size was similar in laser-etched and photolithography-generated devices, while microgels produced in 3D printed devices were generally larger than those generated by the other two techniques, regardless of channel depth (**Figure 5C**). For example, in the 50 µm channel devices, the microgel diameter ranged between ∼ 60-140 µm (in the flow rate ratio of 0.2) depending on the device fabrication technique, with the average microgel diameter being 67.96 ± 9.5 µm for laser etched devices, 61.95 ± 6.68 for photolithography fabricated devices, and significantly higher (p < 0.0001) at 138.71 ± 8.87 µm for 3D printed devices. Moreover, in the same flow rate ratio, the average microgel diameter ranged between ∼ 100-190 μm, and ∼ 160-290μm for channel depth targets of 100 μm, and 400 μm in different techniques.

The diameter of silk microgels was greater when in the oil (before washing in PBS) (**Figure 5E, F**). This characteristic was previously measured and reported as the volumetric swelling ratio (Q_v_ = 0.7) for 3% w/v silk fibroin microgels which were crosslinked with a lower concentration of Ru/SPS of 0.08/0.8 mM [25]. By making silk microgels using different techniques, our results suggest that Q_v_ is independent of the device fabrication method.

Next, the effect of the device fabrication technique on the microgel surface features and their ability to be packed into granular hydrogels was assessed. All fabrication techniques produced spherical microgels with nano-scale pores on the surface (**Figure 6A**), consistent with previous Ru/SPS-crosslinked silk microgels [25] and bulk hydrogels [41]. The overall surface porosity was ∼ 38%, with no significant difference between the porosity of microgels generated using different devices (**Figure 6B**).

Regardless of the fabrication method, all resultant silk microgels could be packed into 3D scaffolds through centrifugation, maintaining structural integrity upon handling (**Figure 6C**). Therefore, despite the 3D printed devices affecting microgel formation efficiency and resulting in larger microgels, all devices successfully produced silk microgels with similar morphological features, that can be used as individual particles or packed into 3D scaffolds.

**Figure 6.**
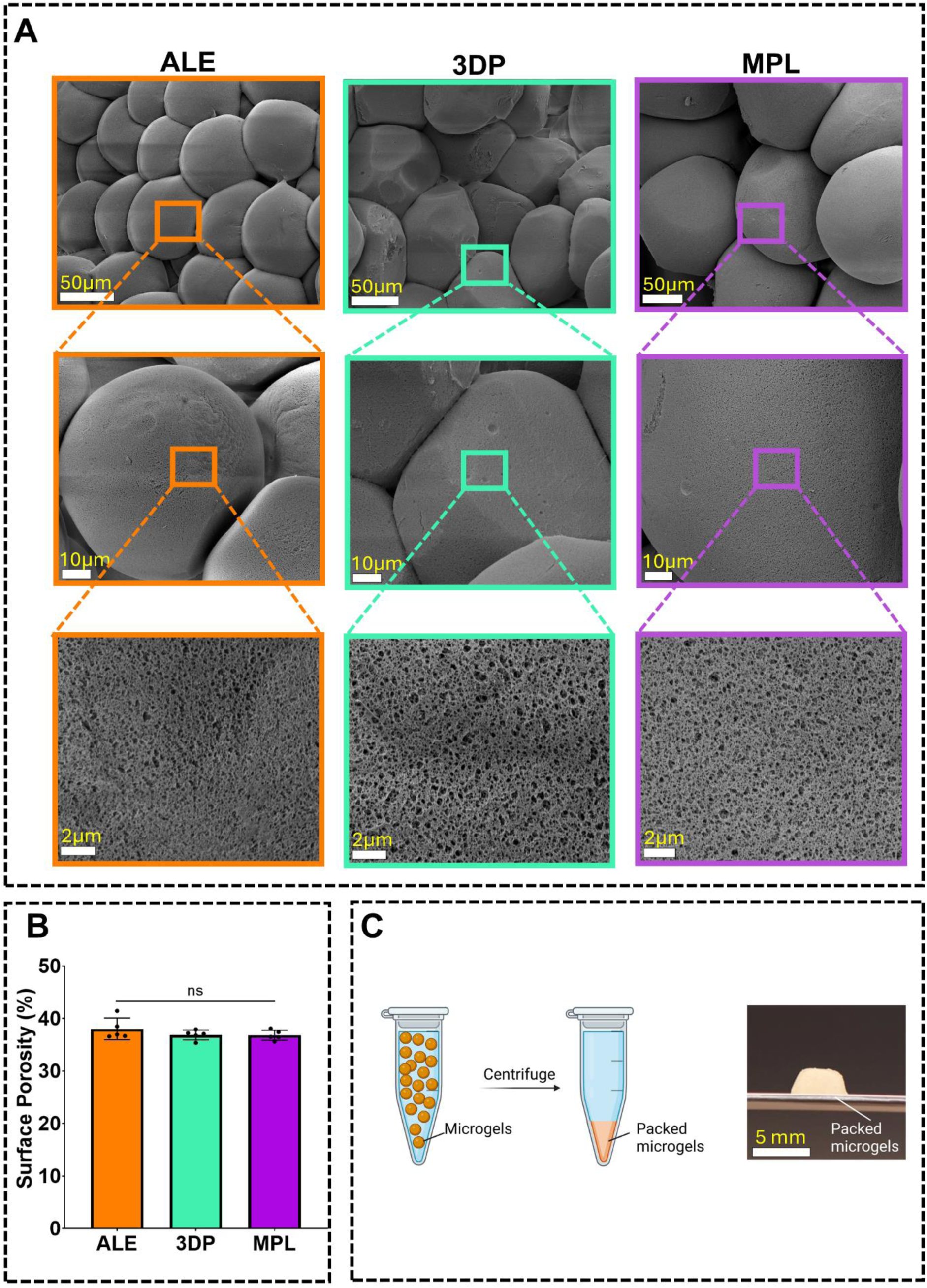
Effect of microfluidic device fabrication technique on the morphological properties of silk microgels. **A)** SEM images of silk microgels made in devices fabricated using three different techniques including acrylic laser engraving (ALE), 3D printing (3DP), and maskless photolithography (MPL). **B)** Surface porosity of silk microgels made using different devices. **C)** Packing silk microgels into a scaffold using centrifugation. Data are mean ± SD, N = 5.

### 3.5. Controlling the size and shape of silk microgels using in-process parameters

Altering the microchannel feature size was shown to affect the size of silk droplets and, consequently, microgels (**Figure 5**). Another strategy for fine-tuning droplet size in passive mode droplet generation is adjusting the flow rate. To explore this, silk microgels were produced using devices made via maskless photolithography, varying the flow rate ratio from 0.1 to 0.5(𝑄_𝑜𝑖𝑙_ = 500 𝜇𝑙/ℎ𝑟) to change the size of microgels formed in the dripping mode (**Figure 7A**). The dripping droplet formation regime was chosen for its high degree of monodispersity and ability to create droplets much smaller than the channel size, avoiding complications associated with fabricating small microfluidic features [40]. For instance, using a microfluidic chip with a channel depth of 400 µm, silk microgels ranging from 210.79 ± 13.72 µm to 411.65 ± 33.15 µm were produced by regulating the silk solution flow rate from 50 𝜇𝑙/ℎ𝑟 to 500 𝜇𝑙/ℎ𝑟 (**Figure 7A**). Overall, silk microgels ranging from 51.11 ± 5.11 µm to 411.65 ± 33.15 µm were successfully fabricated using a combination of channel size and flow rate (**Figure 7A**).

**Figure 7.**
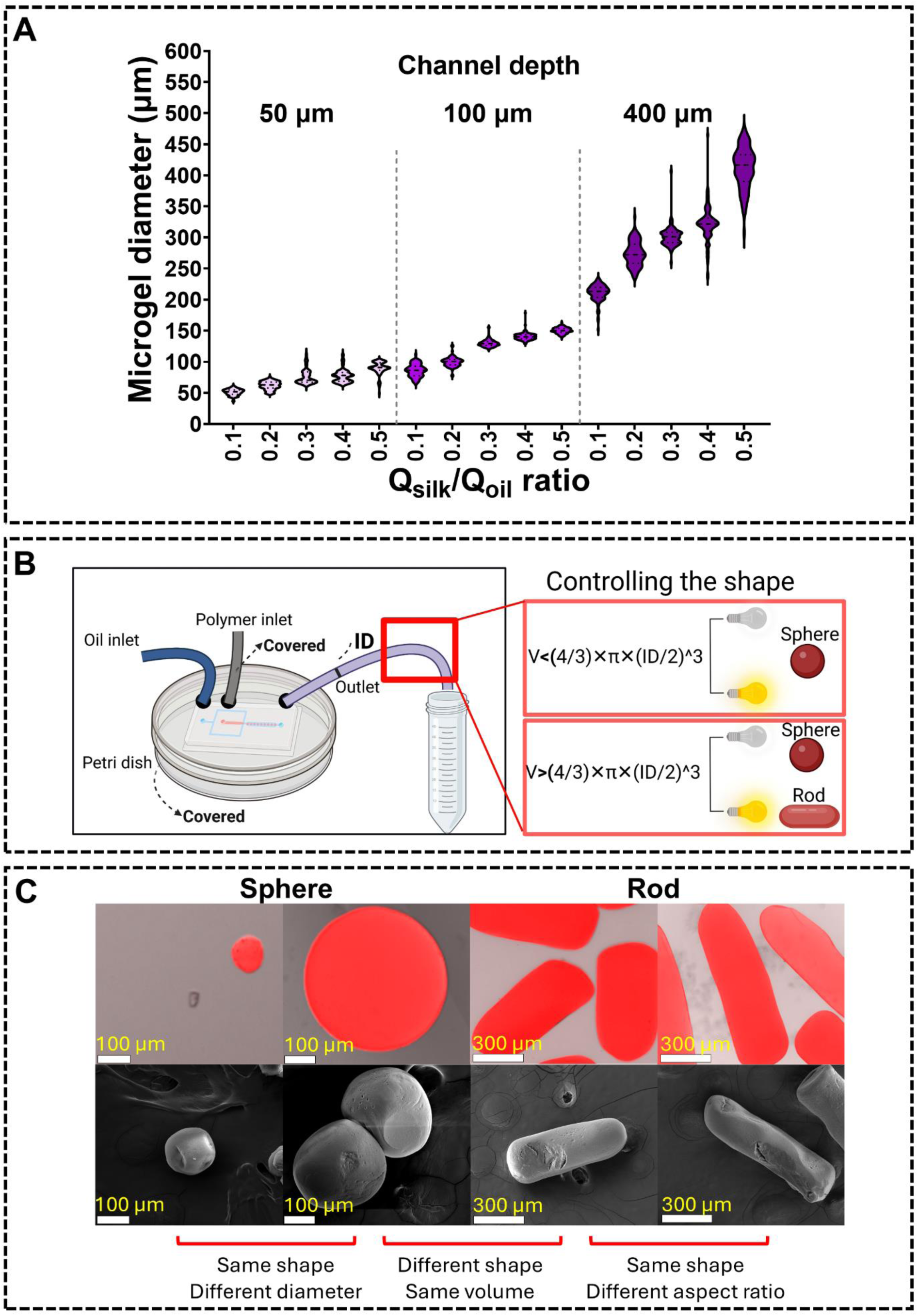
Controlling the size and shape of silk microgels. **A)** Regulating the size of silk microgels by changing the flow rate and microfluidic device channel size. **B)** Regulating in-process parameters to control the shape of the photocrosslinked silk microgels. If the microgels’ diameter is smaller than the inner diameter (ID) of the tubing, they form spheres regardless if they are crosslinked in the tubing or once collected. If the microgel size is larger than the tubing ID, microgels crosslinked in the tubing form rods, while those crosslinked following collection form spheres. **C)** Confocal microscope (top panel) and SEM (bottom panel) images of different silk microgel shapes and sizes generated through the control of in-process parameters. N = 100.

While the microgel size is a function of the channel features and flow rate, the microgel shape is determined by where crosslinking occurs. Photocrosslinking, in particular, allows direct control over microgel shape (**Figure 7B**). For example, if the droplet is large enough (depending on the inner diameter of the tubing), it can be shaped into a sphere or a rod. If droplets are exposed to light while in the outlet tube, they adopt the tube’s shape and form rods. However, if droplets are collected in an oil bath and then crosslinked, they form spherical microgels due to surface tension minimization [42]. By maintaining constant flow rate ratios, microgels with the same volume but different shapes can be produced (**Figure 7C**). This control over the microgel shape and size has important implications for their use in biomedical applications, as these features are known to control drug diffusion from the microgels [43], micropore size when packed in 3D scaffolds [18,25],and biological responses, including cell interactions, tissue infiltration, and immune responses to implanted constructs [25,44].

### 3.6. Effect of silk degumming time (silk molecular weight) on the morphological and mechanical properties of silk microgels

In addition to microgel size and shape, surface morphology and mechanical properties are crucial features that influence biological responses to materials [45,46]. These features are best controlled through silk properties, which are well-documented to affect the final properties of the material [47,48]. Therefore, in this study, the effect of silk molecular weight on the properties of microgels generated in a photolithography-fabricated device with 100 µm channels was examined. Silk molecular weight was controlled by varying the degumming time of silk fibers. While silk degumming results in polydisperse silk populations, longer degumming times produce smaller molecular weight protein species, as well documented in the literature [29,49,50]. The silk degumming time influenced the viscosity of the 3% w/v silk solution used to generate microgels, with viscosity decreasing with longer degumming times (p < 0.01, **Figure 8A**). The viscosity of the 30-min and 60-min (but not 10-min) degummed silk solutions was temperature dependent, decreasing with increased temperature (**Figure 8A**). Silk microgels were successfully generated from all three silk solutions (10-, 30-, and 60-min degummed silk) at a range of flow rates (**Figure 8B**), with microgel size increasing with an increased flow rate in each condition (**Figure 8C**). Despite differences in solution viscosity, the degumming time did not affect microgel size, suggesting that under these conditions the inertial force is dominant over the viscous forces. This is advantageous as it allows making the same size microgels from materials with different viscosities (̴ 2-30 mPa.s) without the need for any further in-process parameters regulation. Moreover, to produce microgels from temperature-sensitive precursors, such as gelatin, alginate, and agarose, where the thermally-induced sol-gel transition is of significance, and viscosity fluctuations may contribute to polydispersity [51], this droplet formation region could be advantageous for maintaining monodispersity.

**Figure 8.**
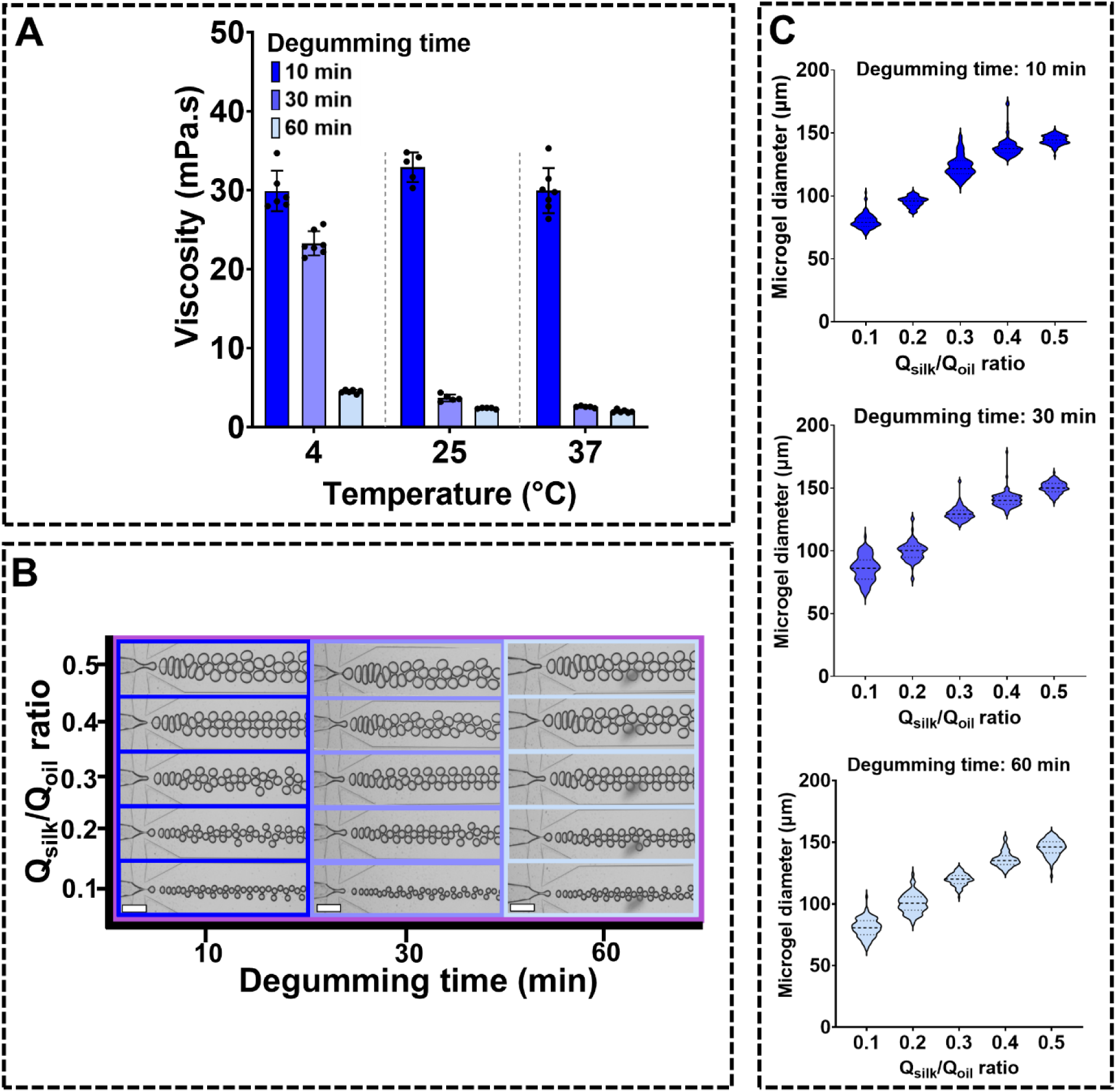
The effect of the silk degumming time on droplet formation and microgels fabrication. **A)** Viscosity of 3% w/v silk solution made from 10-min, 30-min and 60-min degummed silk at different temperatures of 4, 25, and 37°C. **B).** Microfluidic droplet formation from silk with degumming times in various flow rate ratios. **C)** Effect of the degumming time on the size of silk microgel made in different flow rate ratios. All data are from a maskless photolithography-fabricated device with 100 µm channels. N = 100.

While the degumming time did not affect the microgel size, the surface porosity of the silk microgels increased with longer silk degumming times (**Figure 9A**, B). The surface porosity of microgels made from 10-min and 30-min degummed silk was similar at 34.44 ± 1.38% and 38.04 ± 1.85%, respectively, whereas 60-min degummed silk had significantly higher (p < 0.0001, and p < 0.01) surface porosity at 46.34 ± 3.61%. Interestingly, the surface porosity differed from the inner porosity of the microgels (**Figure 9B**), suggesting that either interfacial tension gradients and/or crosslinking gradients affect microgel porosity gradients. This feature should be further explored to control molecular diffusion from the microgels and/or their degradation rates. More generally, the ability to control the morphological properties of silk microgels allows for customization according to specific applications, such as fine-tuning the passive release behavior of molecules for drug delivery systems.

**Figure 9.**
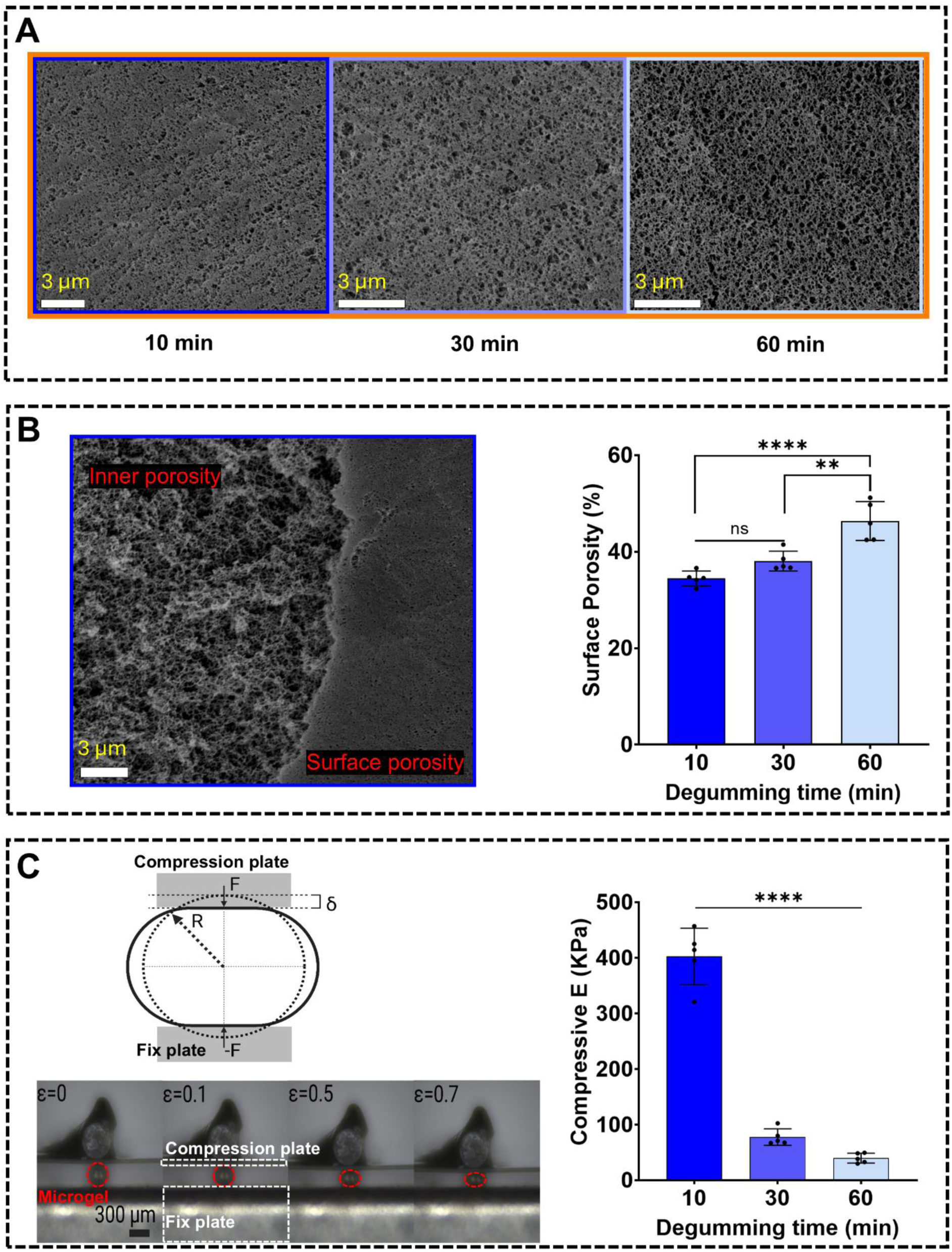
The effect of silk degumming time on the porosity and mechanical properties of silk microgels. **A)** SEM images showing the morphology of silk microgels made using 10-min, 30-min, and 60-min degummed silk solutions. **B)** The effect of degumming time on the surface porosity of silk microgels. **C)** Parallel plate compression testing setup and compressive elastic modulus of silk microgels. All data are from a maskless photolithography-fabricated device with 400 µm channels. Data are mean ± SD, N = 5. ** p < 0.01, **** p <0.0001.

Finally, the mechanical features of single microgels were tested to study the effect of silk degumming time on microgel compression (**Figure 9C**). Increasing degumming time resulted in softer microgels, with those made from 10-min degummed silk having a compressive modulus of 590 ± 144 kPa, significantly higher than that of 30-min degummed silk (77.62 ± 13.29 kPa, p < 0.0001) and 60-min degummed silk (39.81 ± 7.92 kPa, p < 0.0001). When microgels were compressed and unloaded, samples made from 10-min degummed silk exhibited the highest elastic recovery, while 30-min and 60-min degummed samples deformed plastically (viscous energy loss) (**Figure S5B, C**). Therefore, the mechanical properties of silk microgels can be easily tuned for different biomedical applications by changing tfhe silk degumming time while maintaining consistent microgel size.

## 4. Conclusions

This study investigated three common microfluidic device fabrication techniques: acrylic laser engraving, 3D printing with clear resin, and Si-wafer maskless photolithography to produce silk microgels. All three techniques successfully fabricated microfluidic devices with channel depths of 50 µm, 100 µm, and 400 µm. All techniques resulted in over 90% accuracy in channel depth, except for 3D printing, which was not accurate for 50 µm channels. The 3D printed channels were also significantly rougher, resulting in slower microgel production rates across all channel depths, likely due to energy loss and velocity reduction. While maskless photolithography was found to be the most accurate technique, if this specialized method is not available, this research demonstrates the utility of the more accessible and affordable acrylic laser engraving and 3D printing as effective alternatives for generating microgels with similar intrinsic properties.

Silk microgels were fabricated in a wide range of sizes (50 µm to over 400 µm in diameter) by adjusting channel depth and flow rates. Photocrosslinking allowed for precise control over microgel shape, with larger droplets forming rod-like structures when crosslinked in outlet tubing, while post-collection crosslinking produced spherical microgels. Control over the shape and size of microgels is valuable for different applications, including tailoring favorable micro-scale pores and molecular release characteristics.

In addition to microgel size and shape, surface morphology and mechanical properties are crucial for influencing biological responses to materials. This study examined the effect of silk molecular weight, controlled by varying degumming times, on microgels generated in a photolithography-fabricated device. Longer degumming times decreased solution viscosity but did not affect microgel size, allowing consistent microgel production across a range of viscosities (2-30 mPa.s). However, longer degumming times increased surface porosity and decreased compressive modulus, enabling the tuning of mechanical properties for different biomedical applications while maintaining consistent microgel size.

In conclusion, this study successfully demonstrated the versatility and effectiveness of different microfluidic device fabrication techniques for producing photo-crosslinked silk microgels with tunable properties. The ability to control microgel size, shape, surface morphology, and mechanical properties opens numerous possibilities for biomedical applications, including cell encapsulation, cell expansion, drug delivery, and tissue engineering. Future research should focus on exploring annealing strategies to improve injectability for *in vivo* delivery methods, controlling molecular release behavior, and producing more complex structures to further advance the functionality of silk microgels.

## Supporting information

Supporting information is available.

## Acknowledgments

JR-K would like to acknowledge funding support from the Australian Research Council (FT210100668), NSW Health Cardiovascular Research Capacity Program Early-Mid Career Researcher and Collaborative Grants, and UNSW Scientia Program. This work was partly performed at the UNSW Node of the Australian National Fabrication Facility, a company established under the National Collaborative Research Infrastructure Strategy to provide micro and nanofabrication facilities for Australia’s researchers. The authors acknowledge the support from UNSW Engineering Makerspaces and UNSW Design Futures Lab, which provided 3D printing and laser engraving services. The authors thank Dr. Joanna Richmond and Dr. Jake Ireland for technical assistance in SEM imaging carried out at the Electron Microscopy Unit (EMU). EMU is part of the Mark Wainwright Analytical Center at UNSW Sydney, which is in part funded by the Research Infrastructure program at UNSW. The authors thank Mr Hien Tran and Mr Osmond Lao for technical assistance. The schematic drawings were made using Biorender and Onshape. Graphs were generated and statistically analyzed using GraphPad Prism 9 software.

## Contributions

MH, JR-K, NF, and FK conceived the study and designed the experiments. MH carried out experiments, statistical analysis, illustrations, and drafting. All authors aided in interpreting the results, editing, and revising the manuscript.

## References

[1] Drury JL, Mooney DJ. Hydrogels for tissue engineering: Scaffold design variables and applications. Biomaterials 2003;24:4337–51. 10.1016/S0142-9612(03)00340-5.

[2] Zhang YS, Khademhosseini A. Advances in engineering hydrogels. Science (1979) 2017;356. 10.1126/science.aaf3627.

[3] Li J, Mooney DJ. Designing hydrogels for controlled drug delivery. Nat Rev Mater 2016;1. 10.1038/natrevmats.2016.71.

[4] El-Sherbiny IM, Yacoub MH. Hydrogel scaffolds for tissue engineering: Progress and challenges. Glob Cardiol Sci Pract 2013;2013:38. 10.5339/gcsp.2013.38.

[5] Bittner SM, Guo JL, Mikos AG. Spatiotemporal control of growth factors in three-dimensional printed scaffolds. Bioprinting 2018;12. 10.1016/j.bprint.2018.e00032.

[6] Jiménez-Rosado P;, Romero M; Citation: Sánchez-Cid. Polymers (Basel) 2022;2022:3023. 10.3390/polym.

[7] Granular hydrogels for endogenous tissue repair n.d.

[8] Feng Q, Li D, Li Q, Cao X, Dong H. Microgel assembly: Fabrication, characteristics and application in tissue engineering and regenerative medicine. Bioact Mater 2022;9:105–19. 10.1016/j.bioactmat.2021.07.020.

[9] Lai WF, Wong WT. Property-tuneable microgels fabricated by using flow-focusing microfluidic geometry for bioactive agent delivery. Pharmaceutics 2021;13. 10.3390/pharmaceutics13060787.

[10] Dave R, Randhawa G, Kim D, Simpson M, Hoare T. Microgels and Nanogels for the Delivery of Poorly Water-Soluble Drugs. Mol Pharm 2022;19:1704–21. 10.1021/acs.molpharmaceut.1c00967.

[11] Caldwell AS, Aguado BA, Anseth KS. Designing Microgels for Cell Culture and Controlled Assembly of Tissue Microenvironments. Adv Funct Mater 2020;30. 10.1002/adfm.201907670.

[12] de Rutte JM, Koh J, Di Carlo D. Scalable High-Throughput Production of Modular Microgels for In Situ Assembly of Microporous Tissue Scaffolds. Adv Funct Mater 2019;29. 10.1002/adfm.201900071.

13. Granular hydrogels for endogenous tissue repair n.d.

[14] Muir VG, Qazi TH, Shan J, Groll J, Burdick JA. Influence of Microgel Fabrication Technique on Granular Hydrogel Properties. ACS Biomater Sci Eng 2021;7:4269–81. 10.1021/acsbiomaterials.0c01612.

[15] Nguyen TPT, Li F, Hung B, Truong VX, Thissen H, Forsythe JS, et al. Cell Microencapsulation within Gelatin-PEG Microgels Using a Simple Pipet Tip-Based Device. ACS Biomater Sci Eng 2023;9:6024–33. 10.1021/acsbiomaterials.3c00676.

[16] Griffin DR, Weaver WM, Scumpia PO, Di Carlo D, Segura T. Accelerated wound healing by injectable microporous gel scaffolds assembled from annealed building blocks. Nat Mater 2015;14:737–44. 10.1038/nmat4294.

[17] Molley TG, Hung T tyng, Kilian KA. Cell-Laden Gradient Microgel Suspensions for Spatial Control of Differentiation During Biofabrication. Adv Healthc Mater 2022;11. 10.1002/adhm.202201122.

[18] Qazi TH, Wu J, Muir VG, Weintraub S, Gullbrand SE, Lee D, et al. Anisotropic Rod-Shaped Particles Influence Injectable Granular Hydrogel Properties and Cell Invasion. Advanced Materials 2022;34. 10.1002/adma.202109194.

[19] Ou Y, Cao S, Zhang Y, Zhu H, Guo C, Yan W, et al. Bioprinting microporous functional living materials from protein-based core-shell microgels. Nat Commun 2023;14. 10.1038/s41467-022-35140-5.

[20] Zhang P, Zhang Q, Zeng J, Han J, Zhou J, Zhang W, et al. Fabrication of planar photonic crystals in chalcogenide glass film by maskless projection lithography. Appl Phys B 2016;122. 10.1007/s00340-016-6521-x.

[21] Kang M, Han C, Jeon H. Submicrometer-scale pattern generation via maskless digital photolithography. Optica 2020;7:1788. 10.1364/optica.406304.

[22] Thomas T, Agrawal A. Design and fabrication of microfluidic devices: a cost-effective approach for high throughput production. Journal of Micromechanics and Microengineering 2024;34. 10.1088/1361-6439/ad104b.

[23] Nielsen A V, Beauchamp MJ, Nordin GP, Woolley AT. Annual Review of Analytical Chemistry Downloaded from www.annualreviews.org 2020;9. 10.1146/annurev-anchem-091619.

[24] Tweedie M, Maguire PD. Microfluidic ratio metering devices fabricated in PMMA by CO2 laser. Microsystem Technologies 2021;27:47–58. 10.1007/s00542-020-04902-w.

[25] Karimi F, Farbehi N, Ziaee F, Lau K, Monfared M, Kordanovski M, et al. Photocrosslinked Silk Fibroin Microgel Scaffolds for Biomedical Applications. Adv Funct Mater 2024;34. 10.1002/adfm.202313354.

[26] Cai S, Shi H, Li G, Xue Q, Zhao L, Wang F, et al. 3D-printed concentration-controlled microfluidic chip with diffusion mixing pattern for the synthesis of alginate drug delivery microgels. Nanomaterials 2019;9. 10.3390/nano9101451.

[27] Noroozi R, Mashhadi Kashtiban M, Taghvaei H, Zolfagharian A, Bodaghi M. 3D-printed microfluidic droplet generation systems for drug delivery applications. Mater Today Proc, vol. 70, Elsevier Ltd; 2022, p. 443–6. 10.1016/j.matpr.2022.09.363.

[28] Acosta-Cuevas JM, García-Ramírez MA, Hinojosa-Ventura G, Martínez-Gómez ÁJ, Pérez-Luna VH, González-Reynoso O. Surface Roughness Analysis of Microchannels Featuring Microfluidic Devices Fabricated by Three Different Materials and Methods. Coatings 2023;13. 10.3390/coatings13101676.

[29] Rockwood DN, Preda RC, Yücel T, Wang X, Lovett ML, Kaplan DL. Materials fabrication from Bombyx mori silk fibroin. Nat Protoc 2011;6:1612–31. 10.1038/nprot.2011.379.

[30] Nisal A, Sayyad R, Dhavale P, Khude B, Deshpande R, Mapare V, et al. Silk fibroin micro-particle scaffolds with superior compression modulus and slow bioresorption for effective bone regeneration. Sci Rep 2018;8. 10.1038/s41598-018-25643-x.

[31] Bono N, Saroglia G, Marcuzzo S, Giagnorio E, Lauria G, Rosini E, et al. Silk fibroin microgels as a platform for cell microencapsulation. J Mater Sci Mater Med 2023;34. 10.1007/s10856-022-06706-y.

[32] Liu X, Toprakcioglu Z, Dear AJ, Levin A, Ruggeri FS, Taylor CG, et al. Fabrication and Characterization of Reconstituted Silk Microgels for the Storage and Release of Small Molecules. Macromol Rapid Commun 2019;40. 10.1002/marc.201800898.

[33] Toprakcioglu Z, Knowles TPJ. Shear-mediated sol-gel transition of regenerated silk allows the formation of Janus-like microgels. Sci Rep 2021;11. 10.1038/s41598-021-85199-1.

[34] Wiita EG, Toprakcioglu Z, Jayaram AK, Knowles TPJ. Selenium-silk microgels as antifungal and antibacterial agents. Nanoscale Horiz 2024. 10.1039/d3nh00385j.

[35] Green RA, Hassarati RT, Bouchinet L, Lee CS, Cheong GLM, Yu JF, et al. Substrate dependent stability of conducting polymer coatings on medical electrodes. Biomaterials 2012;33:5875–86. 10.1016/j.biomaterials.2012.05.017.

[36] Kim K, Cheng J, Liu Q, Wu XY, Sun Y. Investigation of mechanical properties of soft hydrogel microcapsules in relation to protein delivery using a MEMS force sensor. J Biomed Mater Res A 2010;92:103–13. 10.1002/jbm.a.32338.

[37] Nielsen A V, Beauchamp MJ, Nordin GP, Woolley AT. Annual Review of Analytical Chemistry Downloaded from www.annualreviews.org 2020;9. 10.1146/annurev-anchem-091619.

[38] Amirifar L, Besanjideh M, Nasiri R, Shamloo A, Nasrollahi F, De Barros NR, et al. Droplet-based microfluidics in biomedical applications. Biofabrication 2022;14. 10.1088/1758-5090/ac39a9.

[39] Sattari A, Hanafizadeh P, Hoorfar M. Multiphase flow in microfluidics: From droplets and bubbles to the encapsulated structures. Adv Colloid Interface Sci 2020;282. 10.1016/j.cis.2020.102208.

[40] Nunes JK, Tsai SSH, Wan J, Stone HA. Dripping and jetting in microfluidic multiphase flows applied to particle and fibre synthesis. J Phys D Appl Phys 2013;46. 10.1088/0022-3727/46/11/114002.

[41] Tran HA, Maraldo A, Ho TTP, Thai MT, van Hilst Q, Joukhdar H, et al. Probing the Interplay of Protein Self-Assembly and Covalent Bond Formation in Photo-Crosslinked Silk Fibroin Hydrogels. Small 2024. 10.1002/smll.202407923.

[42] Fitzpatrick R. Surface tension. Theoretical Fluid Mechanics, IOP Publishing; 2017, p. 3-1–3–21. 10.1088/978-0-7503-1554-8ch3.

[43] Adepu S, Ramakrishna S. Controlled drug delivery systems: Current status and future directions. Molecules 2021;26. 10.3390/molecules26195905.

[44] Veiseh O, Doloff JC, Ma M, Vegas AJ, Tam HH, Bader AR, et al. Size- and shape-dependent foreign body immune response to materials implanted in rodents and non-human primates. Nat Mater 2015;14:643–51. 10.1038/nmat4290.

[45] Shin JW, Jang JY, Lee SW, Park SH, Shin JW, Mun C, et al. Combined effects of surface morphology and mechanical straining magnitudes on the differentiation of mesenchymal stem cells without using biochemical reagents. J Biomed Biotechnol 2011;2011. 10.1155/2011/860652.

[46] Oluwole SA, Weldu WD, Jayaraman K, Barnard KA, Agatemor C. Design Principles for Immunomodulatory Biomaterials. ACS Appl Bio Mater 2024. 10.1021/acsabm.4c00537.

[47] Lin Y, Xia X, Shang K, Elia R, Huang W, Cebe P, et al. Tuning chemical and physical cross-links in silk electrogels for morphological analysis and mechanical reinforcement. Biomacromolecules 2013;14:2629–35. 10.1021/bm4004892.

[48] Johari N, Moroni L, Samadikuchaksaraei A. Tuning the conformation and mechanical properties of silk fibroin hydrogels. Eur Polym J 2020;134. 10.1016/j.eurpolymj.2020.109842.

[49] Wray LS, Hu X, Gallego J, Georgakoudi I, Omenetto FG, Schmidt D, et al. Effect of processing on silk-based biomaterials: Reproducibility and biocompatibility. J Biomed Mater Res B Appl Biomater 2011;99 B:89–101. 10.1002/jbm.b.31875.

[50] Rnjak-Kovacina J, Wray LS, Burke KA, Torregrosa T, Golinski JM, Huang W, et al. Lyophilized Silk Sponges: A Versatile Biomaterial Platform for Soft Tissue Engineering. ACS Biomater Sci Eng 2015;1:260–70. 10.1021/ab500149p.

[51] Jiang W, Li M, Chen Z, Leong KW. Cell-laden microfluidic microgels for tissue regeneration. Lab Chip 2016;16:4482–506. 10.1039/c6lc01193d.

