## Supplemental information for "Evaluating Flow-Focused Microfluidic Device Fabrication Techniques for Silk Fibroin Microgel Production"

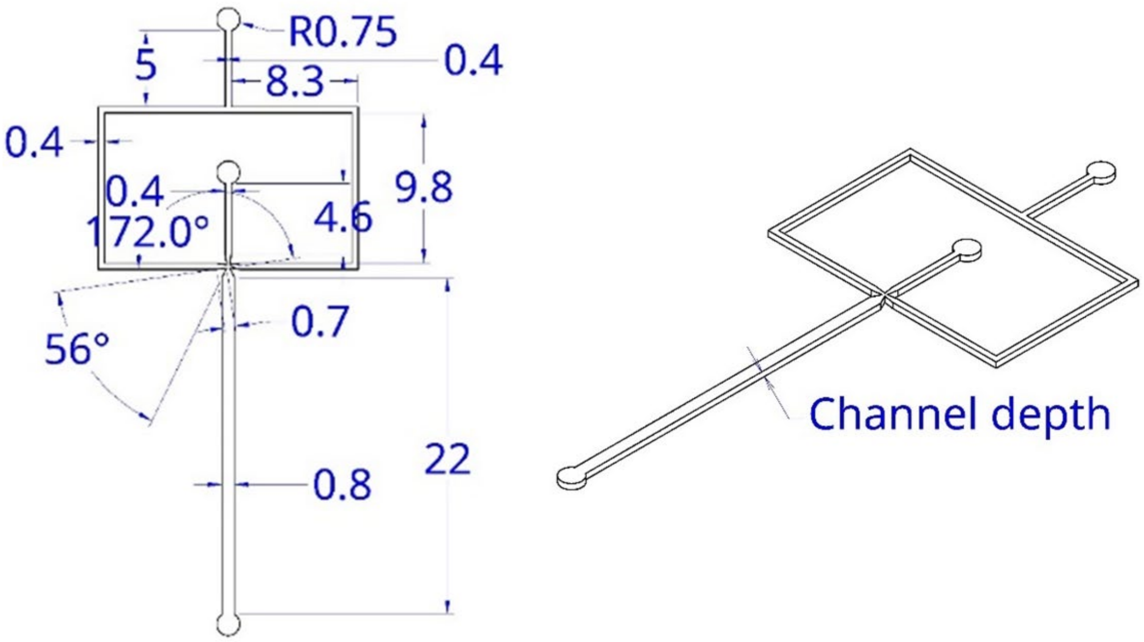


***Figure S1. Microfluidic device channel design.***

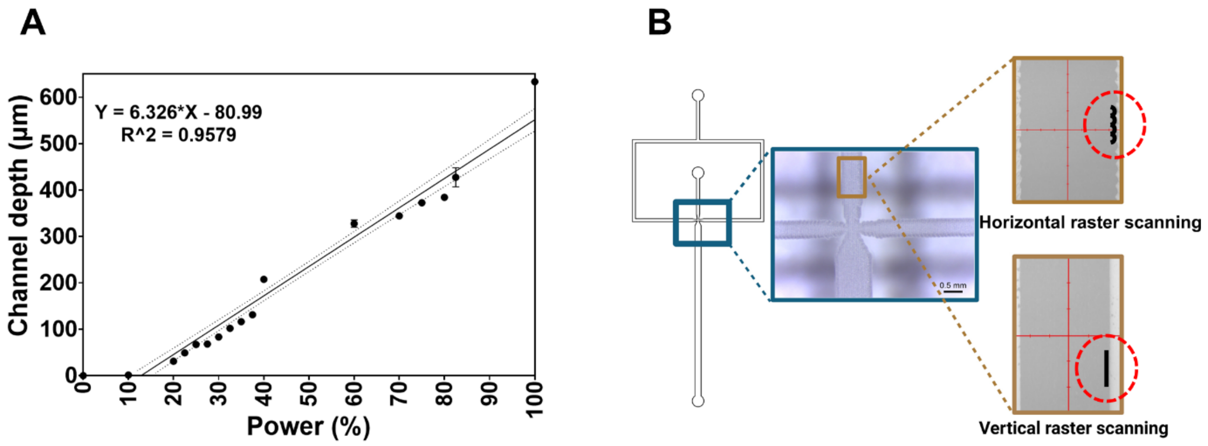


***Figure S2. Optimizing $CO_2$-assisted laser engraving of the acrylic substrate. A)*** *laser power-engraving depth relationship.* ***B)*** *Effect of raster scanning direction on channel patterning. N = 3-5.*

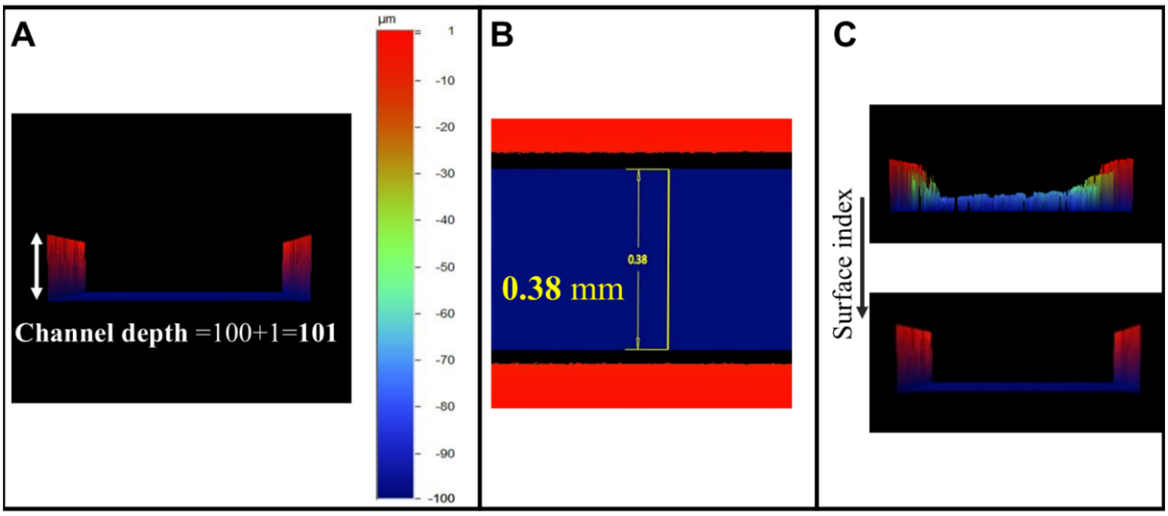


***Figure S3. Characterizing the channel geometry using optical profilometry. A)** Channel depth was measured as the peak-to-valley height value (Z-axis). **B)** Channel width measurement (X-axis). **C)** Surface index measurement as an indicator of surface roughness.*

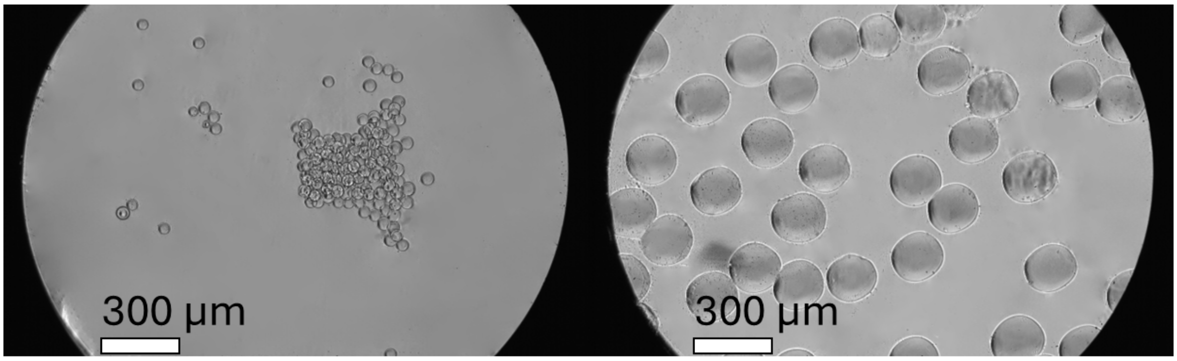


***Figure S4. Silk microgel morphology under an optical microscope.***

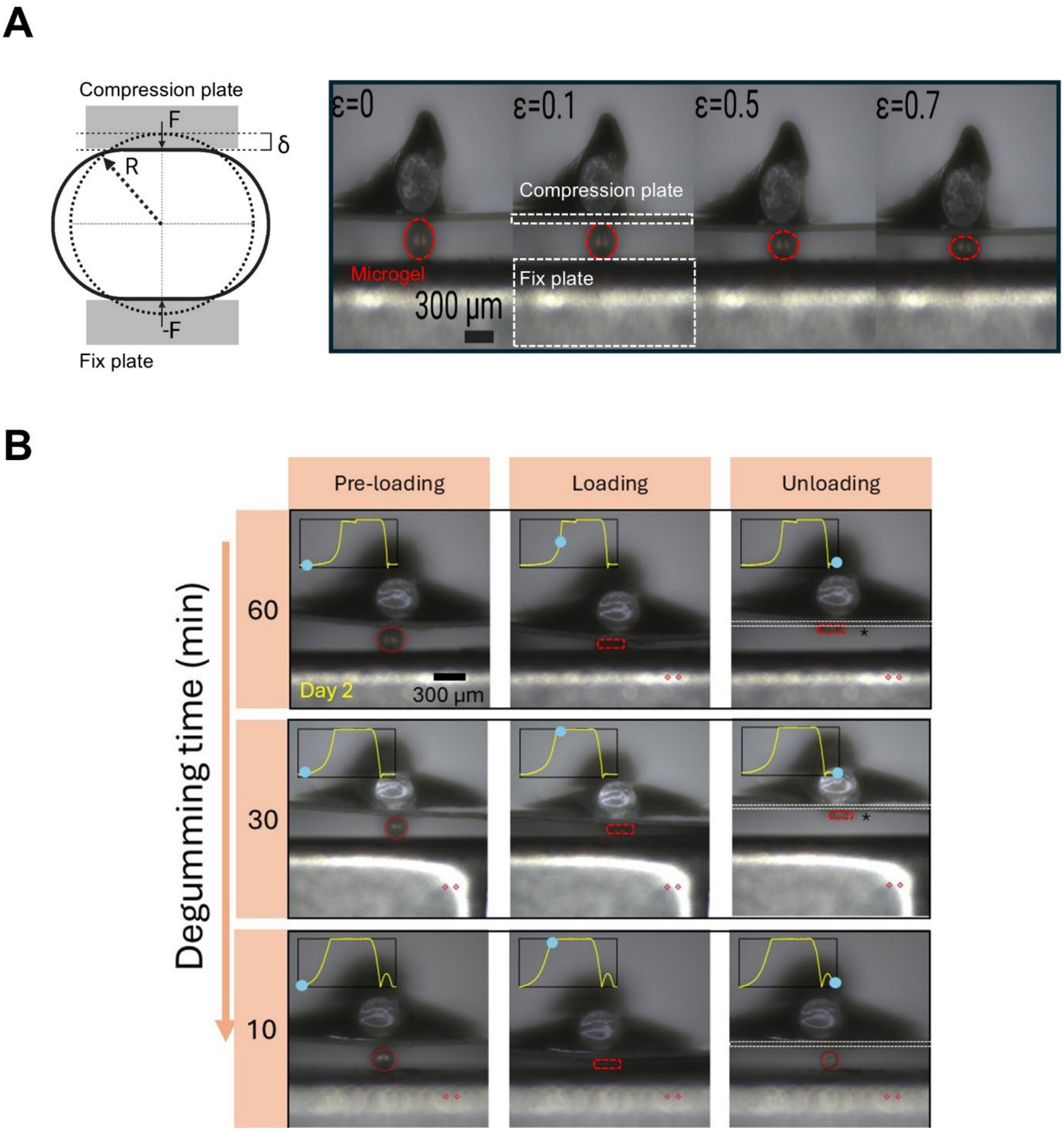


***Figure S5. Parallel plate compression testing of single microgels. A)*** *Using the Hertzian half-space contact model and parallel plate compression testing of microgels to calculate the compressive elastic modulus.* ***B) Silk microgels on day 2 before, while, and after deformation (200 µm displacement magnitude during the 20s).***
